# A kinetic model improves off-target predictions and reveals the physical basis of *Sp*Cas9 fidelity

**DOI:** 10.1101/2020.05.21.108613

**Authors:** Behrouz Eslami-Mossallam, Misha Klein, Constantijn v.d. Smagt, Koen v.d. Sanden, Stephen K. Jones, John A. Hawkins, Ilya J. Finkelstein, Martin Depken

**Affiliations:** Kavli Institute of NanoScience and Department of BioNanoScience, Delft University of Technology, Delft 2629HZ, the Netherlands; Department of Molecular Biosciences, University of Texas at Austin, Austin, Texas 78712, USA; Institute for Cellular and Molecular Biology, University of Texas at Austin, Austin, Texas 78712, USA; Center for Systems and Synthetic Biology, University of Texas at Austin, Austin, Texas 78712, USA; Oden Institute for Computational Engineering and Science, University of Texas at Austin, Austin, Texas 78712, USA

**Author notes:** equal contribution (Behrouz Eslami-Mossallam, Misha Klein).

## Abstract

The *S. pyogenes (Sp)* Cas9 endonuclease is an important gene-editing tool. *Sp*Cas9 is directed to target sites via a single guide RNA (sgRNA). However, SpCas9 also binds and cleaves genomic off-target sites that are partially matched to the sgRNA. Here, we report a microscopic kinetic model that simultaneously captures binding and cleavage dynamics for *Sp*Cas9 and *Sp*-dCas9 in free-energy terms. This model not only outperforms state-of-the-art off-target prediction tools, but also details how *Sp-*Cas9’s structure-function relation manifests itself in binding and cleavage dynamics. Based on the biophysical parameters we extract, our model predicts *Sp*Cas9’s open, intermediate, and closed complex configurations and indicates that R-loop progression is tightly coupled with structural changes in the targeting complex. We show that *Sp*Cas9 targeting kinetics are tuned for extended sequence specificity while maintaining on-target efficiency. Our extensible approach can characterize any CRISPR-Cas nuclease – benchmarking natural and future high-fidelity variants against *Sp*Cas9; elucidating determinants of CRISPR fidelity; and revealing pathways to increased specificity and efficiency in engineered systems.

---

CRISPR-Cas9 (Clustered Regularly Interspaced Short Palindromic Repeats – CRISPR associated protein 9) is a ubiquitous tool in the biological sciences^1,2^ with applications ranging from live-cell imaging^3^ and gene knockdown/overexpression^4,5^, to genetic engineering^6,7^ and gene therapy^8,9^. *Streptococcus pyogenes* (*Sp*) Cas9 is programmed with a ∼100 nucleotide (nt) single-guide RNA (sgRNA) to target DNAs based on the level of complementarity to a 20 nt segment of the sgRNA^10^. Wild type *Sp*Cas9 (Cas9 from now on) induces specific double-stranded breaks and the catalytically ‘dead’ Cas9 (dCas9) mutants allow for binding the target DNA without cleavage^3,5^. Apart from complimentary *on-targets*, Cas9-sgRNA also binds and cleaves partially-complementary *off-target* DNA sites ^11–18^. Off-target cleavage risks unwanted genomic alterations, including point mutations, large-scale deletions, and chromosomal rearrangements^19^. The potentially deleterious effects associated with such editing errors impedes wide-spread implementation of the CRISPR toolkit in human therapeutics.

Off-target sites are identified *in silico* by a growing set of prediction tools. These tools use bioinformatics^20,21^, machine learning^22,23^, and heuristic^12,14,24,25^ approaches to rank genomic sites based on their own unique off-target activity scores. However, none of these tools attempt to model the microscopic kinetic properties that govern Cas9-DNA binding and nuclease activation. This quantitative kinetic modeling is essential for understanding how *in vivo* Cas9 activity depends on enzyme concentration and exposure time. Both of these parameters are frequently exploited by experimentalists to limit off-target activity in cells^26^.

Quantitative predictions of Cas9 activity requires a physical model that accounts for the kinetic nature of the problem. Existing physical models^24,27^ implicitly assume that Cas9-sgRNA binding equilibrium is reached over the entire genome before DNA cleavage. However, binding does not necessarily equilibrate before cleavage^28,29^, as can be inferred from the fact that binding and cleavage correlate weakly *in vitro* and in cells^30–32^ (see below). Here, we construct a comprehensive kinetic model that includes binding and cleavage reactions, and globally train it on two high-throughput *in vitro* datasets that capture each process separately^15^. Our fully parameterized model accurately predicts an independent high-throughput dataset^11^, without the use of any additional fitting parameters. Our model is parameterized in terms of physical quantities and therefore offers insights into biophysical mechanisms. By establishing the free-energy landscape of the targeting reaction with any off-target, which shows that the difference in binding and cleavage activities^30–39^ stems from a (relatively) long-lived DNA-bound intermediate. We further show that this state is tuned for both high cleavage specificity and on-target cleavage efficiency. We also connect the binding intermediate to the intermediate HNH-conformation observed in single-molecule FRET experiments^40,41^, and argue that the conformational change is driven by R-loop formation. Finally, we show that our kinetic model outperforms the two best-performing genomic off-target prediction tools used today^12,24,42^.

## Results

### Kinetic model simultaneously captures binding and cleavage profiles

**Figure 1a** describes the microscopic kinetic schema that underpins our physical Cas9 binding and cleavage model. First, the Cas9-sgRNA ribonucleoprotein complex recognizes a 3nt protospacer adjacent motif (PAM) DNA sequence—canonically 5’-NGG-3’—via protein-DNA interactions^43,44^. Binding to the PAM sequence opens the DNA double helix, and allows the first base of the target sequence to hybridize with the sgRNA^43,44^. The DNA double helix further denatures as the sgRNA and target strand form an RNA-DNA hybrid (R-loop)^45–48^. The R-loop grows and shrinks in single nucleotide steps until it is either reversed and Cas9 dissociates, or it reaches completion (a 20 nt hybrid). If the R-loop reaches completion, Cas9 uses its HNH and RuvC nuclease domains to cleave both strands of the DNA duplex^49^.

**Fig. 1.**
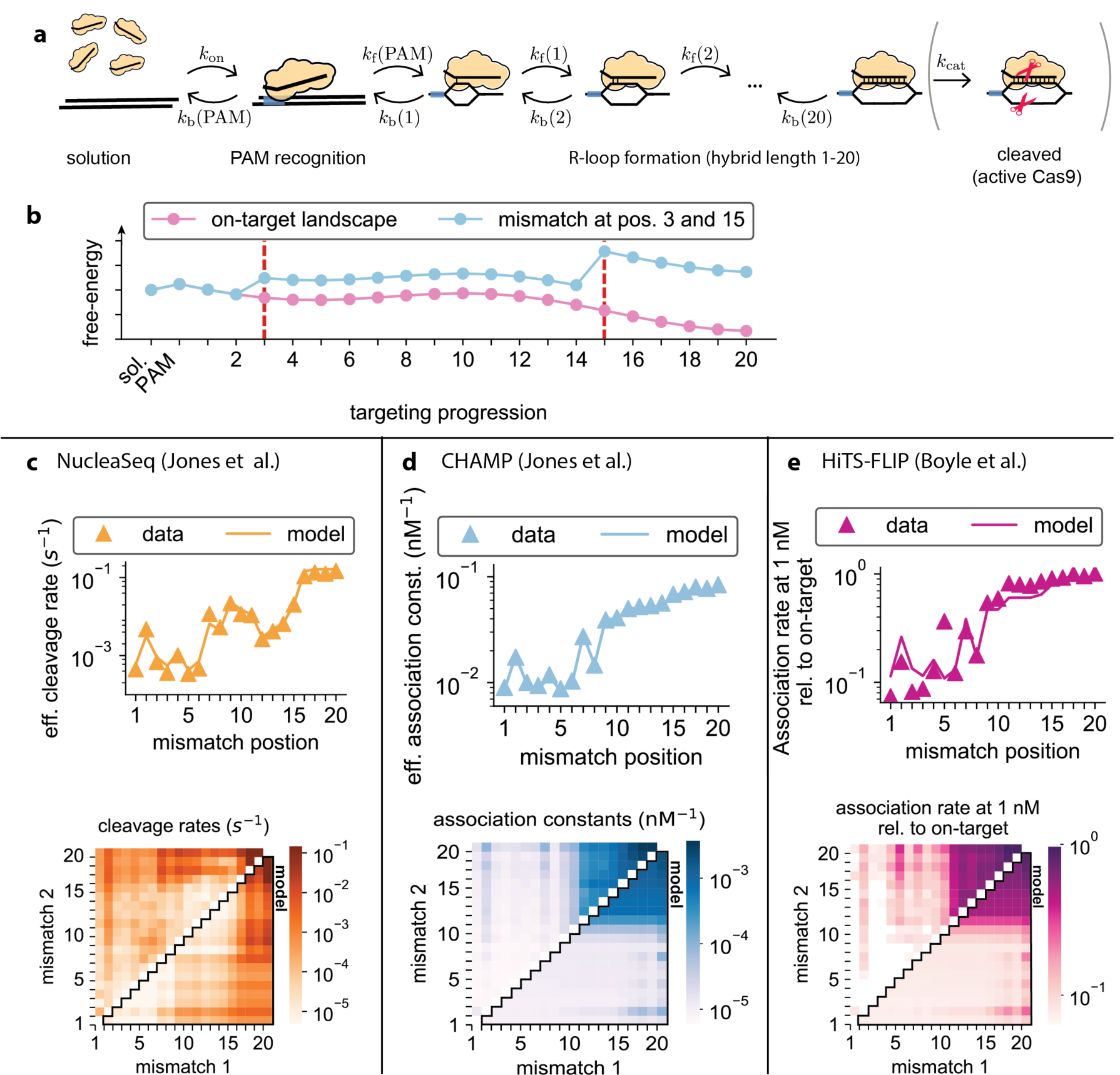
A kinetic model captures both binding and cleavage data. **a**, Reaction schema underlying the proposed kinetic model (**Supplementary Information** for details). An available (d)Cas9-sgRNA from solution binds a DNA sequence (either on- or off-target) at its PAM site (blue rectangle) with rate *k*_on_. R-loop formation then proceeds in one base-pair increments. A partially formed R-loop containing *n* base pairs can either extend one base pair at a rate *k*_f_(*n*) or shrink one base pair at a rate *k*_b_(*n*). A complete R-loop (20 base pairs) is cleavage competent, and a dsDNA break is catalyzed at a rate *k*_cat_. For dCas9, cleavage catalysis is not available, and *k*_cat_ = 0. **b**, Illustration of a possible free-energy landscape for Cas9-sgRNA-DNA for the on-target (pink) and an off-target with mismatches placed at positions 3 and 15 (blue). Each mismatch raises the entire free-energy landscape starting from the position where it occurs. **c**, Effective cleavage rates and **d**, effective association constants as measured by and simultaneously fitted (**Supplementary Information**) to the NucleaSeq and CHAMP datasets (*Jones at el*.). Off-targets with one mismatch are show on top and off-targets with two mismatches are shown at the bottom (data above and model below diagonal), both as a function of mismatch position(s). **e**, Model prediction for effective binding rates as a function of mismatch position(s) compared to both HiTS-FLIP data and data from *Boyle et al*. top: one mismatch; bottom: two mismatches, with experimental data above and model results below the diagonal.

The most general reaction schemes for cleavage and binding are completely parameterized only when we estimate all the rates shown in **Fig. 1a** for every potential guide-target combination—a prohibitively large number of parameters for any genome. To render parameter estimation tractable, we make four mechanistic model assumptions: (1) mismatch positions within the hybrid are more important than mismatch types—*e*.*g*. G-G vs. G-A (as can be inferred directly from data^11,15^), and all 12 mismatch types can be treated equally; (2) dCas9 differs from Cas9 only in that dsDNA bond-cleavage catalysis is completely suppressed, and all other rates can be taken to be identical between the two^40,50^; (3) a mismatch influences only the reversal of the mismatched base pairing, leaving all other rates unchanged; (4) all hybrid-bond-formation rates are equal, and independent of complementarity. These assumptions are justified *post hoc* by showing that the targeting dynamics are completely determined by even a much smaller set of effective rates. Though our model is kinetic, we can use the detailed-balance condition for microscopic rates (**Supplementary Information**) to define the free-energy of each state in our model (**Fig. 1b**). Our model assumptions reduce the total number of parameters to 44: the (concentration dependent) rate of PAM binding from solution (*k*_on_) and the associated free-energy cost; a single internal forward (bond-formation) rate (*k*_f_); 20 free-energy costs dictating R-loop progression for matching guide and target; 20 free-energy penalties for mismatches at different R-loop positions; and, for Cas9, the rate at which the final cleavage reaction is catalyzed (*k*_cat_) (see **Supplementary Information** for further details). When extending the R-loop, both gains and losses in free-energy are possible as base-pairing interactions, protein-DNA interactions^50^, and any induced conformational changes^40,41,49,51^ all contribute to the stability of the Cas9-sgRNA-DNA complex. As we assume that mismatches only facilitate the reversal of the mismatched base pairs, the entire free-energy landscape will rise by a positive amount from the mismatch onwards (c.f. pink and blue free-energy landscapes in **Fig. 1b**).

We used three high-throughput assays to train and validate our kinetic model. The first training data set estimates the effective cleavage rates (*k*_clv_) for a library of off-target DNA sequences by monitoring the fraction of uncut DNA over time^15^ (NucleaSeq in **Fig. 1c**; **Supplementary Information**). The second training data set reports on the effective association constant (*K*_A_) over a library of off-target DNA sequences exposed to dCas9-sgRNA for 10 min^15,52^ (CHAMP in **Fig. 1d; Supplementary Information**). The third data set, used for validating the model, reports the effective association rate estimated over 1500 seconds of exposure to dCas9-sgRNA at 1nM concentration (HiTS-FLIP in **Fig. 1e**; **Supplementary Information**)^11^. Our kinetic model describes all these experiments even though each dataset uses either Cas9 or dCas9 to report on the cleavage rates, association constants, or association rates by sweeping either concentration or time^28^.

We trained the kinetic model on DNA binding (CHAMP) and cleavage (NucleaSeq) datasets collected using the same sgRNA and mismatched target DNA library^15^. The parameters were globally fit to the binding affinities and cleavage rates for all off-target DNA sequences with up to two mismatches. The rates from different types of mismatches were averaged together (**Supplementary Information**). Although these two datasets do not correlate well with each other directly (55%, see **Supplementary Figure 1a**), our model reproduces effective cleavage rates (**Fig. 1c**) and effective association constants (**Fig. 1d**) with a high correlation (86% and 99%, respectively; **Supplementary Figures 1b** and **c**). As validation, our model accurately captures a third, independent dataset of dCas9 effective association rates^11^ (**Fig. 1e**) with a correlation of 97% (**Supplementary Figure 1d**), and without the use of additional fitting parameters. Our model also predicts the CHAMP data for sequences with more than 2 mismatches, even though these were not included in the training data (**Supplementary Figures 1e** and **f**). We conclude that our model accurately captures the physics of DNA binding and cleavage by Cas9.

### Internal R-loop states are tuned for cleavage specificity without loss of on-target efficiency

To gain mechanistic insights into the targeting reactions, we investigated the estimated free-energy landscape and kinetic parameters (**Fig. 2**) resulting from the simultaneous fit to our training datasets **Figs. 1c,d**). Starting from the PAM-bound state, the on-target free-energy (**Fig. 2a**) increases substantially when forming the first hybrid base pair and remains relatively high until the 8^th^ base pair is formed. This initial barrier must be bypassed before a stable binding intermediate is reached with about 11 or 12 hybridized base pairs. The free-energy landscape reveals a second barrier to forming a full R-loop (13-18 bp), and eventual cleavage. The penalty for a mismatch (**Fig. 2b**) contains contributions from both DNA-RNA base pairing and protein-nucleotide interactions. Still, the mismatch penalties remain rather constant throughout (6±1 *k*_B_*T*), with notable exceptions being positions 2, 18 and 20.

**Fig. 2.**
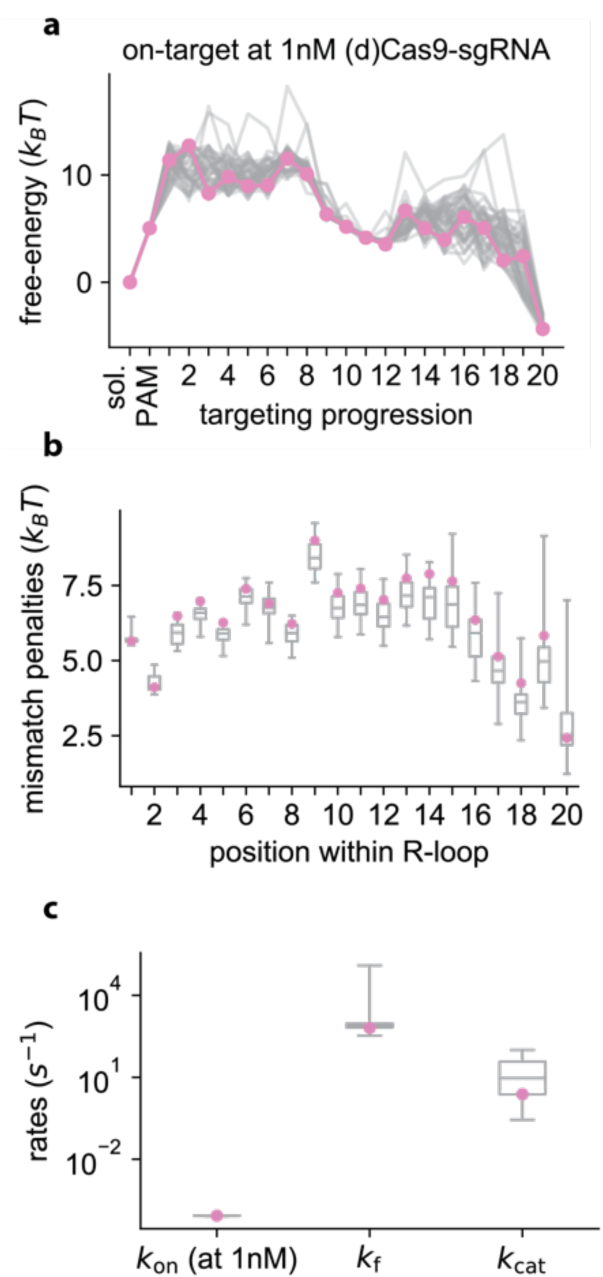
Microscopic parameter estimated from NucleaSeq and CHAMP datasets. **a**, The free-energy landscape of the on-target reaction along the states shown in **Fig. 1a**. Here sol. is the solution state, PAM is the PAM-bound state and numbers indicate the number of R-loop base pairs formed. **b**, Energetic penalties for mismatches as a function of position. **c**, The three forward rates. In all panels, the global fit with the lowest chi-squared is shown in pink (**Supplementary Information**), and all nearly optimal solutions are represented in grey. For the lower two panels, the interquartile range of nearly optimal solutions are represented in the grey boxes and whiskers denote the complete range of values.

If the system equilibrates between major barriers in the free-energy landscape, we expect that any change in barrier height can be compensated for by the appropriate change in the bond-formation rate (*k*_f_) (**Supplementary Figures 2a**,**b**)—without effecting model predictions. Consequently, both quantities cannot be simultaneously determined in a partially equilibrated system, explaining the high variability of predicted barrier heights (**Fig. 2a**) and *k*_f_ (**Fig. 2c**). By directly showing that the predicted binding and cleavage profiles are indeed insensitive to changing the barrier height (**Supplementary Figures 2c** and **d**), as long as the forward rate is appropriately adjusted, we confirm partial equilibration of the system. This insight both explains the high variance of free-energy estimates in barrier regions (**Fig. 2a**), and allows us to perform coarse-grain modeling of the system to isolate parameters that are well determined by the data.

Based on the free-energy landscapes in **Fig. 2a**, we identified equilibrated states as those with free energies that are well-constrained by the fits. The equilibrated states are the effective states used in our coarse-grained model, and we calculate the coarse grained parameter values based on the estimated parameter values of the full model (**Supplementary Information**). We define the open (O) R-loop state as the PAM bound state. The local minimum in **Fig. 2a** defines our coarse-grained intermediate (I) R-loop state with between 7 and 13 of its hybrid base pairs formed. Finally, the closed (C) R-loop and cleavage-competent state contains a fully formed hybrid. The resulting coarse-grained reaction scheme (**Fig. 3a**) captures the experimental data as well as the complete model (**Supplementary Figure 3**). This coarse-grained model reveals the rate-limiting steps during on- and off-target DNA binding and cleavage.

**Fig. 3.**
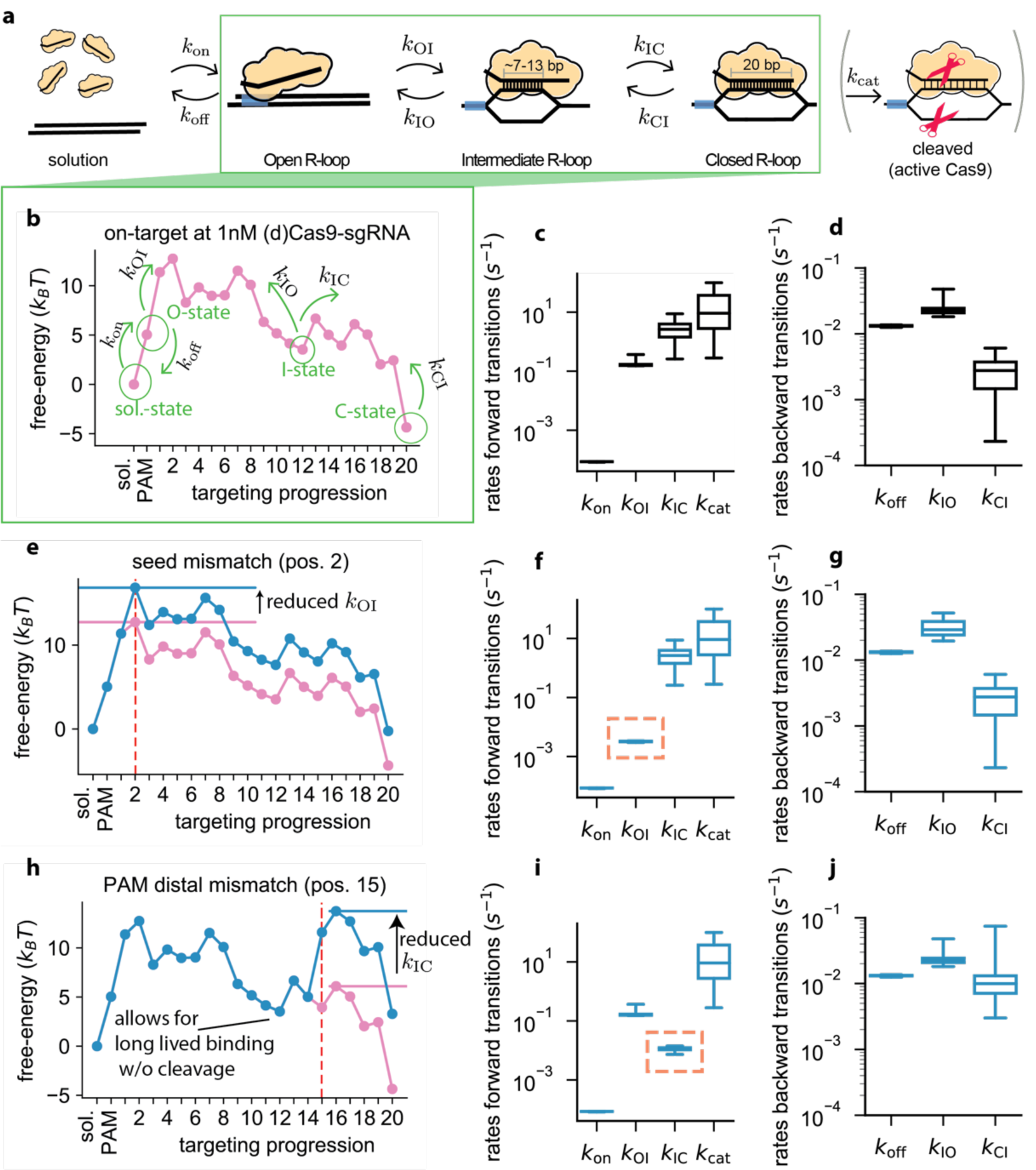
Coarse-grained kinetic model fully captures bulk data. **a**, Coarse-grained version of the reaction scheme shown in **Fig. 1a**. Apart from the unbound and post-cleavage state, the targeting-reaction pathway is reduced to just three states (open, intermediate, and closed R-loops, see **Supplementary Information** for details). **b**, Microscopic free-energy landscape for the on-target exposed to 1nM (d)Cas9-sgRNA (**Fig. 2a**) with coarse-grained states and rates indicated in green. **c**, Coarse-grained forward and **d**, backward rates associated with the landscape in **b. e**, Microscopic free-energy landscape for an off-target with a mismatch at position 2 exposed to 1nM (d)Cas9-sgRNA (blue), together with the on-target free-energy landscape (pink). **f, g**, Coarse-grained forward (**f**) and backward (**g**) rates associated with the landscape in **e. h-j**, Same as (**f-g**) for an off-target with a mismatch at position 15.

The rate-limiting step for on-target cleavage is the transition from the open to the intermediate R-loop state (*k*_OI_ ≪ *k*_IC_) (**Figs. 3b-d**). Complexes that enter the intermediate state also typically enter the closed state (*k*_IO_ ≪ *k*_IC_). The transition between the open and the intermediate state is reversible because the free-energy difference between the open and intermediate state is low (resulting in *k*_IO_ ≈ *k*_OI_). The free-energy difference between the intermediate and closed state is high (*k*_IC_ ≫ *k*_CI_), rendering the transition from an opened to closed configuration essentially irreversible and all but guarantee cleavage (*k*_CI_ ≪ *k*_cat_).

Mismatches between the target DNA and the sgRNA have differential effects on R-loop propagation. A PAM-proximal seed mismatch (R-loop nucleotides 1-8) suppresses the rate of transition from an open to intermediate state 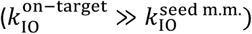 (**Figs. 3e-g**). In contrast, a PAM-distal mismatch (R-loop nucleotides 12-17) limits the effective rate of cleavage from the open state by reducing the intermediate to closed state transition (*k*_IC_ ≪ *k*_OI_) (**Figs. 3h-j**). The transition from binding to the intermediate state remains unaffected, though returning to the open state competes with completion of the R-loop (*k*_IO_ ≈ *k*_IC_). Enzymes that enter the closed state likely also proceed to cleavage (*k*_CI_ ≪ *k*_cat_). We conclude that specificity in PAM-distal regions (i.e., the second barrier in off-target landscape shown in **Fig. 3h** is higher than the first) is tuned not to interfere with the crucial on-target cleavage efficiency (i.e., second barrier in **Fig. 3b** is lower than the first).

### R-loop propagation drives Cas9 conformation dynamics

What are the structural properties of Cas9 that give rise to the non-monotonic free-energy landscape of **Fig. 2a**? A comparisons between DNA-bound and unbound Cas9-sgRNA structures have revealed that Cas9 repositions HNH and RuvC nuclease domains to catalyze cleavage^44,53,54^. We hypothesized that the position of the mobile HNH nuclease domain directly couples to R-loop progression, allowing it to influence its free-energy landscape. This hypothesis is based on the following key observations: First, ensemble FRET experiments^49^ detected two dominant Cas9 conformers with distinct HNH states, and single-molecule FRET studies have identified a third intermediate conformer^40,41,51^—matching the number of R-loop states we find; Second, the relative position and occupancy of the HNH states is affected by R-loop mismatches^40,41,51^, while the Cas9 can only sense mismatches by hybridizing the sgRNA with the target DNA.

To test this hypothesis, we mimicked the experiments of Dagdas *et al*.^40^ by calculating the time evolution of the occupancy for each of the microscopic states in the DNA-bound Cas9 landscape for three target sequences (**Fig. 4**). The HNH-domain completes its conformational change within seconds after Cas9-sgRNA binds to on-target DNA^40^. Our model demonstrates a similar behavior for R-loop progression (**Fig. 4a**). The intermediate R-loop state (green) is visited only transiently, while the closed state (red) strongly resists unwinding of the full hybrid (*k*_IC_ ≫ *k*_CI_) (**Figs. 3c, d** and **4a**). Compared to the on-target DNA, PAM-distal mismatches reduce the intermediate closure rate (*k*_IC_) and increase the time spent in the intermediate state (**Fig. 4b**), in agreement with prior observations^40^. Our model also shows how going from three to four PAM distal mismatches effectively abolishes the occupancy of the closed state at short times^40^, as R-loop formation is stalled in the intermediate state (**Fig. 4c**). In prior FRET experiments, the FRET value corresponding to the intermediate state depended on the number of mismatches introduced, which is evidence that the HNH domain adopts slightly different configurations^41^. The reported relationship between FRET values and mismatches is consistent with tight coupling of conformational change to R-loop progression in PAM distal regions, as our model predicts that going from three to four PAM-distal mismatches increases the probability of residing as a larger intermediate R-loop (**Fig. 4**). Taken together, we propose that the three coarse-grained R-loop states identified in our free-energy landscape reflect the three HNH domain conformers. The free-energy landscape (**Fig. 2** and **3**) obtained by fitting bulk data (**Fig. 1**) thus complements structural and single-molecule data to describe how Cas9 targets matched and mismatched DNAs.

**Fig. 4.**
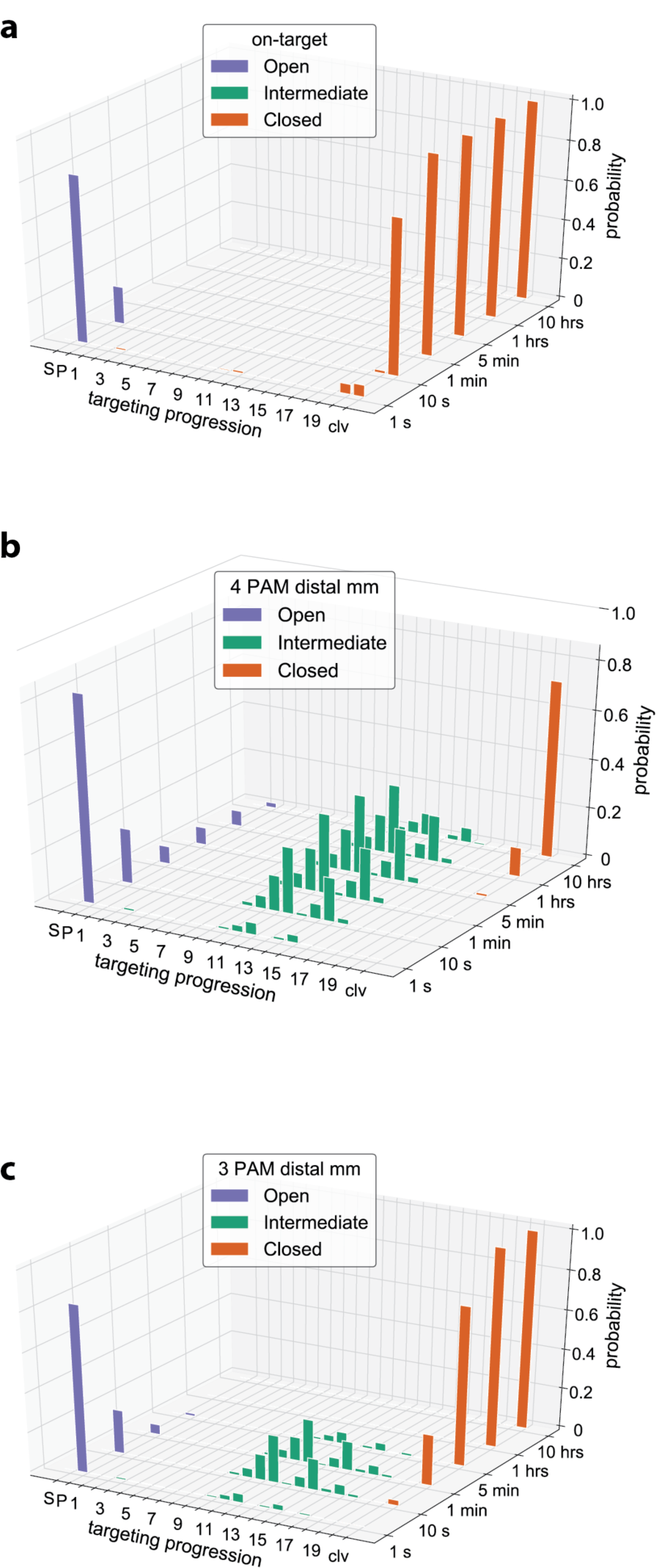
The time evolution of R-loop hybridization is reminiscent of conformational dynamics. **a**, The evolution of the occupation probability for any of the 23 microscopic states shown in **Fig. 1a**, as a function of time when interacting with the on-target. **b**, Same as a but when interacting with an off-target with last (PAM distal) 3 base pairs mismatched. **c**, Same as a but when interacting with an off-target with the last 4 base pairs mismatched. Colors indicate the corresponding coarse-grained R-loop configuration as defined in **Fig. 3a**: open R-loop and unbound states (blue), intermediate R-loop states (green) and cleavage-competent and post-cleavage states (orange).

### Kinetic modelling improves genome-wide off-target prediction

Next, we sought to exploit our mechanistic description for predictive power, and compared our predictions with those of current state-of-the-art genomic off-target prediction tools. Current methods^12,14,20,21,23–25,42^ rank genomic off-targets according to various measures of *in vivo* activity without predicting biochemically measurable parameters (i.e., the binding affinity or cleavage rate). One such frequently-used tool computes the Cutting Frequency Determination (CFD) score^12^ —a naïve-Bayes classification scheme^22^ that assumes mismatches affect the relative cleavage probabilities multiplicatively. More recently, Zhang *et al*. presented a unified CRISPR (uCRISPR) score that outperforms the CFD score^24^. uCRISPR estimates the cleavage probability as proportional to the Boltzmann weight corresponding to the cleavage competent state. The assumption of a multiplicative effect errors and the use of Boltzmann weights both implicitly imply binding equilibrium. This assumption is not borne out by the experimental data as the off-target binding and cleavage patterns do not match (see e.g. **Supplementary Figure 1a**).

To analyze whether our kinetic model improves genomic off-target predictions, we collected data from sequencing-based cleavage experiments. To comprehensively evaluate the model, we gathered data from all experiments that used industry-standard sgRNAs (i.e., targeting EMX1, FANCF, HBB, RNF2, and VEGFA) and reported multiple off-target cleavage sites^33–36,38,39^. Notably, these reports identified only partially overlapping sets of off-target cleavage sites, indicating that off-target cleavage detection is strongly dependent on experimental parameters (i.e., Cas9 nucleofection vs. plasmid transfection, exposure time, cell type, etc.) and the sensitivity of detection (i.e., enrichment of breaks or whole-genome sequencing)^16,18^. For each sgRNA, we separately tested against the union (sites found in any experiment) and intersection (sites found in every experiment) of the reported off-target sites (**Fig. 5** and **Supplementary Figures 4-6**). The union of all reported off-targets maximizes the likelihood of covering low probability off-targets, while the intersection minimizes the effect of experiment-dependent biases and noise.

**Fig. 5.**
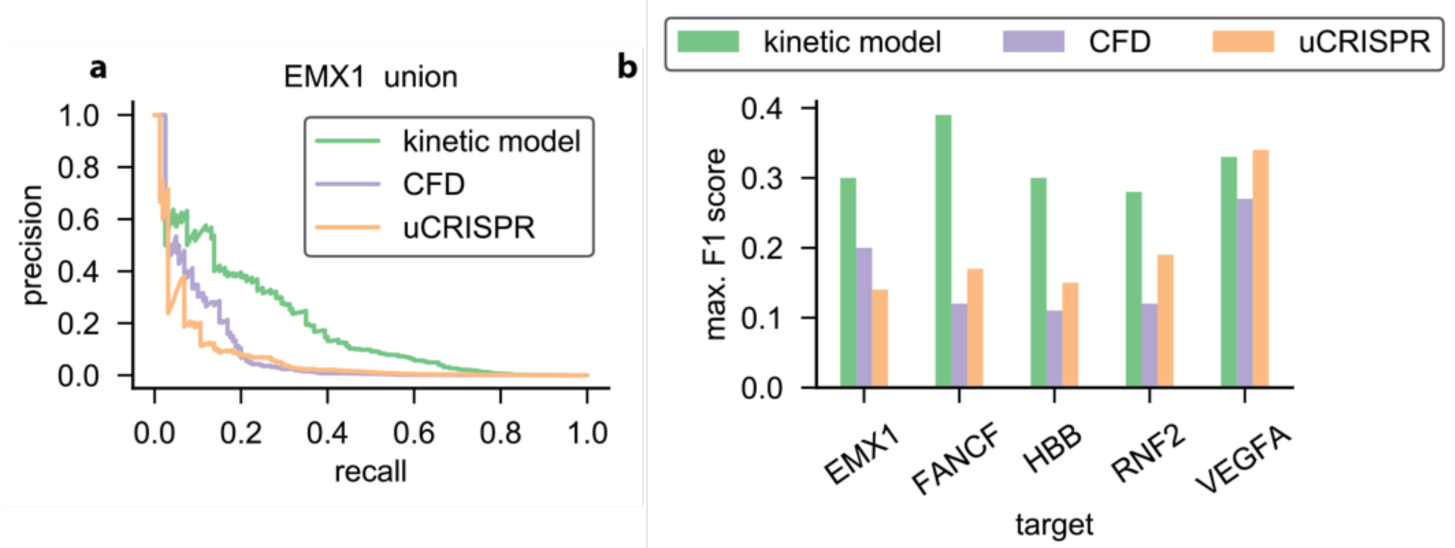
Genome-wide off-target classification. **a**, Precision recall-curves for our model (green) and predictions based on the CFD score (purple) or uCRISPR score (orange) for the EMX1 site using all experimentally identified off-targets. **b**, F1-scores for our model (green), CFD prediction tool (purple) and uCRISPR (orange), for target sites EMX1, FANCF, HBB, RNF2 and VEGFA site 1 using all experimentally identified off-targets. For each condition, the maximum obtainable F1-score along the corresponding PR-curve is displayed (see **a** and **Supplementary Figure 4**)

We tested how well our model, the CFD score, and uCRISPR separate reported off-targets over the human genome. For sake of comparison, we need to collapse our dynamic description into a binary classification. We choose to separate out strong off-targets based on the predicted cleavage vs. unbinding probability once the Cas9-sgRNA has bound the PAM^28^, as this is proportional to the steady-state cleavage rate in the low concentration limit. To simplify the comparison further, we only considered sequences flanked by a canonical NGG motif. **Fig. 5a** shows the resulting precision-recall (PR) curve when tested against all reported off-targets of the EMX1 guide sequence (union). As the threshold for strong off-targets is swept, PR curves display the fraction of sites that are correctly labelled as off-target (precision) against the fraction of the experimentally-identified sequences that are predicted (recall). For therapeutic genome-editing, a high recall is imperative as a false negative prediction is more harmful than a false positive one. Our kinetic model produces higher recall values for all achievable precisions, clearly outperforming state-of-the-art CFD and uCRISPR classifying schemes for EMX1 (**Fig. 5a**). Importantly, the kinetic model also outperforms the leading off-target predictors for highly-mismatched genomic off-targets of other sgRNAs, as judged by PR-curves, receiver operating characteristic curves, and the F1-score (**Fig. 5** and **Supplementary Figures 4-6**). This result is especially surprising since the kinetic model was trained on datasets that captured, at most, two mismatches from a single on-target sequence.

## Discussion

Here, we describe a kinetic model for Cas9 binding and cleavage that is trained on high-throughput *in vitro* measurements^15^. This bottom-up modelling approach has allowed us to decipher the microscopic free-energy landscape underlying *Sp*Cas9 target recognition (**Fig. 1-2**). Based on extracted free-energy landscapes, we find that *Sp*Cas9’s kinetics are dominated by transitions between the open, intermediate, and closed R-loop states (**Fig. 3**). As mismatches affect the three R-loop states similarly to the three configurational states of Cas9’s nuclease domains^40,41^, we propose that PAM distal R-loop formation is tightly coupled to protein conformation (**Fig. 4**)—pointing toward the relevant structure-function relation for the most important RNA-guided nuclease in use today.

By mechanistically accounting for the kinetic nature of the targeting process, our model outperforms existing genome-wide off-target prediction tools. For simplicity and robustness, we built our model to exclude mismatch type parameters, allowing for extensive training using datasets based on a single guide sequence and off-target DNAs containing up to two mismatches. This training does not limit the model’s application as the model also improves on the detection of highly-mismatched genomic off-target sites (**Fig. 5** and **Supplementary Figures 4-6**).

Our model is also the first to fully capture the time dependence of off-target binding in addition to cleavage. Understanding the time dependence of off-target binding will facilitate the design of sgRNA libraries in Cas9- or dCas9-based experiments. For example, a recent study by Jost *et al*.^5^ demonstrated that a series of mismatched guides can be used to titrate gene expression during CRISPRa/CRISPRi. Knowing *Sp*Cas9’s microscopic free-energy landscape (**Figs. 2-3**) can also simplify the design of CRISPRa/CRISPRi libraries for novel gene targets. Wildtype Cas9 can also be (effectively) inactivated with PAM-distal mismatches in the guide^55^, and our model can guide titration of Cas9-sgRNA inactivation.

The physical insights generated by the free-energy landscapes we extract could also help rational protein-reengineering efforts aimed at producing high-fidelity Cas9 variants that maintain high on-target efficiency^39,51,56^. For *Sp*Cas9, we find that the barrier between the intermediate and closed states is tuned to extend the cleavage specificity beyond the seed, without affecting on-target efficiency (**Figs. 3b** and **h**).

Taken together, we have shown that mechanistic modelling combined with high-throughput data sets give biophysical insights into *Sp*Cas9 off-targeting, and that those insights give predictive power far beyond the training sets. *Sp*Cas9 is only one of many RNA-guided nucleases with biotechnological applications, and other CRISPR associated nucleases (such as Cas12a, Cas13 and Cas14) offer a diversified genome-engineering toolkit^15,57–62^. These nucleases can all be characterized with our approach, and it will be especially interesting to compare the free-energy landscape of our *Sp*Cas9 benchmark to that of engineered^39,51,56^ and natural (e.g. *N. meningitides* Cas9^63^) high-fidelity Cas9 variants.

## Acknowledgements

We would like to thank Kristian Blom, Diewertje Dekker, and Sonny de Jong for valuable discussions and their help during the project. Thank you also to the members of the Chirlmin Joo lab and Stan Brouns lab for valuable discussions. We also thank Evan Boyle for sharing his data and answering all our questions. B.E.M. forms part of the research program ‘‘Crowd management: the physics of genome processing in complex environments”, supported by NWO. M.K. was supported by the Netherlands Organization for Scientific Research (NWO/OCW), as part of the Frontiers in Nanoscience program. M.D. acknowledges support from the Parents in KIND program, sponsored by The Kavli Institute of Nanoscience Delft, the Department of Bionanoscience at TU Delft, and through a Spinoza Prize awarded to M. Dogterom by NWO. I.J.F. is supported by a University of Texas College of Natural Sciences Catalyst award and the Welch Foundation (F-1808). I.J.F. and S.K.J. are supported by the U.S. National Institute of Health (R01GM124141, F32AG053051).

## Author contributions

B.E.M and M.K: designed and performed the research. Wrote the manuscript

K.v.d.S and C.v.d.S: Performed the research.

S.K.J: Provided data. Wrote manuscript

J.H: Provided data. Wrote manuscript

I.J.F: Provided data. Wrote manuscript

M.D: Designed the research. Wrote manuscript

