## Supplementary Information for "A kinetic model improves off-target predictions and reveals the physical basis of *Sp*Cas9 fidelity"

### 1 Modeling targeting kinetics

We here explain the details of the model introduced in the main text (**Fig.** 1a). We consider a single DNA target sequence with PAM, in contact with a Cas9-sgRNA solution at fixed concentration. After a Cas9-sgRNA binds the DNA at its PAM site with a rate proportional to the Cas9-sgRNA concentration, R-loop formation (Cas9 mediated strand exchange between gRNA and DNA) is modeled as a sequential process where the sgRNA-DNA hybrid stochastically grows and shrinks with single-nucleotide increments. The DNA strand can only be cleaved once a complete R-loop is formed (20 bp). We model target recognition as a hopping process along the sequence $\mathrm{sol}, \mathrm{PAM}, 1, 2, ...., 20, \mathrm{clv}$ of states. The process starts in the sol state, where our DNA molecule is empty, and eventually ends in the absorbing clv state, where the DNA molecule is cleaved. We will for notational convenience also refer to the same sequence of states as $-1, 0, 1, 2, ...., 20, 21$.

#### 1.1 The Master equation and its general solution

Letting $P_n(t)$ denote the probability to be in state $n$ at time $t$, and $k_{\mathrm{f}}(n)/k_{\mathrm{b}}(n)$ the rates for forward $(n \to n+1)$/backward $(n \to n-1)$ transitions, we can describe the evolution of the probabilities with the Master equation

$$\frac{\partial P_{-1}}{\partial t} = -k_{\mathrm{f}}(\text{-}1)P_{-1}(t) + k_{\mathrm{b}}(0)P_0(t) \tag{S1}$$

$$\cdots$$

$$\begin{aligned}\frac{\partial P_n}{\partial t} &= k_{\mathrm{f}}(n-1)P_{n-1}(t) - (k_{\mathrm{f}}(n) + k_{\mathrm{b}}(n))P_n(t) \\ &+ k_{\mathrm{b}}(n+1)P_{n+1}(t)\end{aligned} \tag{S2}$$

$$\cdots$$

$$\frac{\partial P_{20}}{\partial t} = k_{\mathrm{f}}(19)P_{19}(t) - (k_{\mathrm{f}}(20) + k_{\mathrm{b}}(20))P_{20}(t) \tag{S3}$$

The fraction of cleaved DNA (for active Cas9) is directly set by the fraction of uncleaved DNA, $P_{21}(t) = 1 - \sum_{n \leq 20} P_n(t)$, and we do not need to explicitly include the cleaved (21st) state in the Master equation. Defining the vector $\vec{P}(t) = [P_{-1}(t),\, P_0(t),\, P_1(t), \ldots, P_{20}(t)]^T$, the formal solution to **Equations** S1 and S2 can be written as

$$\vec{P}(t) = e^{-Kt}\vec{P}(0), \tag{S4}$$

where $K$ is a tri-diagonal rate matrix. If we define $k_{\mathrm{f}}(-1) = k_{\mathrm{b}}(21) = 0$, we can give the elements of $K$ as

$$K_{nm} = \begin{cases} -k_{\mathrm{f}}(n-1) & n = m+1 \\ k_{\mathrm{f}}(n) + k_{\mathrm{b}}(n) & n = m \\ -k_{\mathrm{b}}(n+1) & n = m-1 \\ 0 & |n-m| > 1 \end{cases}, \quad n, m \in \{-1, 0, 1, \ldots, 20\} \tag{S5}$$

#### 1.2 The mechanistic model assumptions

In the text we introduce four mechanistic model assumptions: (1) we can average over mismatch types, and only keep track of matches and mismatches; (2) dCas9 has all the same rates as Cas9, apart from that the cleavage rate $k_{\mathrm{cat}} = k_{\mathrm{f}}(20) = 0$. (3) Introducing a mismatch at $n$, changes only $k_{\mathrm{b}}(n)$; (4) $k_{\mathrm{f}}(0) = k_{\mathrm{f}}(1) = \ldots = k_{\mathrm{f}}(19) \equiv k_{\mathrm{f}}$, and independent of mismatch pattern.

We use the reference concentration 1nM ([Cas9-sgRNA] $= C_{\mathrm{Cas9-sgRNA}}/1\mathrm{nM}$), and take the rate of binding from solution to grow linearly with concentration $k_{\mathrm{f}}(-1) = k_{\mathrm{on}}^{1\mathrm{nM}}$[Cas9-sgRNA]. Taken together, forward transitions are assigned the following rates

$$k_{\rm f}(n) = \begin{cases} k_{\rm on}^{\rm 1nM}[\text{Cas9-sgRNA}], & n=-1, \\ k_{\rm f}, & n \in [0,1,...,19], \\ k_{\rm cat}, & n=20. \end{cases} \tag{S6}$$

##### 1.3 Parameterization in terms of free energies

The collection of all backward and forward rates completely determined the probabilistic evolution by the system, and thus calculate the ensemble average of any quantity defined over our state space. Still, it is informative to translate these rates into free-energy differences between states. To do this, we imagine that two neighboring states to be allowed to mutually equilibrate. Introducing the free energy of state $n$ as $F_n$ (in units of $k_{\rm B}T$), equilibration between states $n$ and $n+1$ means both that the relative occupancy is described by Boltzmann weights $P_n^{\rm EQ}/P_{n+1}^{\rm EQ} = \exp[-(F_n - F_{n+1})]$ and that there are no net probability currents between the states $P_n^{\rm EQ}k_{\rm f}(n) = P_{n+1}^{\rm EQ}k_{\rm b}(n+1)$. Taken together, these relationships tie rates to free-energy differences through

$$F_n - F_{n-1} = \ln[k_{\rm b}(n)/k_{\rm f}(n-1)]. \tag{S7}$$

This equation, together with model assumption (1)-(4) and our assumption of first order binding to the PAM, can be used to write

$$F_n - F_{n-1} = \begin{cases} F_{\rm PAM}^{\rm 1nM} - \ln([\text{Cas9-sgRNA}]), & n=0 \\ \epsilon_{\rm C}(n), & \text{match at } n \in [1,2,...20] \\ \epsilon_{\rm C}(n) + \epsilon_{\rm I}(n) & \text{mismatch at } n \in [1,2,...20]. \end{cases} \tag{S8}$$

If the $n^{\rm th}$ base of the target is complementary to the corresponding base of the guide, the energy of the Cas9-sgRNA-DNA ternary complex increases by $\epsilon_{\rm C}(n) \times k_B T$ when incorporating the basepair into the R-loop. The Cas9 protein is known to interact with both DNA strands, as well as undergo conformational changes during the process of R-loop formation, and $\epsilon_{\rm C}(n)$ can take both positive ($k_{\rm f}(n-1) < k_{\rm b}(n)$) and negative ($k_{\rm f}(n-1) > k_{\rm b}(n)$) values. If the $n^{\rm th}$ base of the target does not match the guide's base, the ternary complex gets penalized with an energetic cost $\epsilon_{\rm I}(n) \geq 0$.

In conclusion, we have built a general mechanistic model in terms of rates and **Equation** S4, and have re-parametrized it in terms of 41 free energies ($F_{\rm PAM}^{\rm 1nM}$, $20 \times \epsilon_{\rm C}(n)$'s, $20 \times \epsilon_{\rm I}(n)$) and three forward rates ($k_{\rm on}^{\rm 1nM}$, $k_{\rm f}$, and $k_{\rm cat}$).

#### 2 Calculating measured quantities

Here we explain how we calculated quantities to compare to experimental high-throughput data sets for training and validation.

##### 2.1 Calculating effective cleavage rates for NucleaSeq

We use the solution to the Master Equation (**Equation** S4) to calculate the expected cleaved fraction for Cas9 at any complementarity pattern, and compare this to the read counts acquired during the NucleaSeq experiment [1]. NucleaSeq is performed at saturating concentrations of Cas9-sgRNA, which we model by setting $F_{\rm PAM} = F_0 = -1000$. As done in the original experiment, we record the fraction of DNA that remains uncleaved ($\sum_{n\leq 20} P_n(t)$) at the time points 0 s, 12 s, 60 s, 180 s, 600 s, 1800 s, 6000 s, 18000 s, and 60000 s, and fit out a single effective cleavage rate $k_{\rm clv}$.

#### 2.2 Calculating effective association constants for CHAMP

We model the CHAMP experiments [1] by calculating the bound fraction ($P_{\text{bnd}}(t) = \sum_{n=0}^{20} P_n(t)$) of dCas9 ($k_{\text{f}}(20) = k_{\text{cat}} = 0$) after 10 min at concentrations 0.1 nM, 0.3 nM, 1 nM, 3 nM, 10 nM, 30 nM, 100 nM and 300 nM. Assuming the system has had sufficient time to equilibrate within the 10 minutes for which the experiment lasts, the series of occupancies should follow the Hill equation

$$P_{\text{bnd}}^{\text{EQ}} = \frac{[\text{Cas9-sgRNA}]}{[\text{Cas9-sgRNA}] + 1/K_{\text{A}}}. \tag{S9}$$

We follow the experimental protocol and fit **Equation** S9 to the bound fraction to extract apparent association constant $K_{\text{A}}$ for the (off-)target of interest.

#### 2.3 Calculating effective association rates for HiTS-FLIP

To predict measured association rates in the HiTS-FLIP experiment [2], we again compared the flourecence signal to our calculated bound fraction ($P_{\text{bnd}}(t) = \sum_{n=0}^{20} P_n(t)$) for dCas9. Following Boyle et al. [2] we use linear regression to fit a straight line passing through the origin to the bound fraction ($P_{\text{bnd}} \approx k_{\text{on}}^{\text{eff}} t$) at time points $t_1 = 500s$, $t_2 = 1000s$ and $t_3 = 1500s$.

### 3 Parameter estimation

Here we describe the global cost function we seek to minimize, as well as the algorithms used to search for the global minimum.

#### 3.1 A global cost function

All fits are performed using a custom written Simulated Annealing (SA) algorithm [3] (see below) to minimize the total relative square devitation between data and prediction. The global cost function $\chi^2$ used for training adds contributions from the NucleaSeq data and the CHAMP data,

$$\chi^2 = \chi^2_{\log_{10} k_{\text{clv}}} + \chi^2_{\log_{10} K_{\text{A}}} \tag{S10}$$

As the recorded rates and association constants span many orders of magnitude, we will seek to minimize the relative square deviation; we do this by minimizing the square deviation of the log measures $\log_{10}(k_{\text{clv}})$ or $\log_{10}(K_A)$. The response of $k_{\text{clv}}$ and $K_{\text{A}}$ show clear patterns for both one and two mismatches, while especially the cleavage activity is much lower for three mismatches and above. We therefor limit our training set to one and two mismatches. The number of available mismatch configurations with two mismatches ($^{20\times19}/_{2} = 190$ in total) exceeds that of single mismatches (20 in total), and to equalize the weight of each data set we consider the total residues averaged over their respective number of configurations

$$\chi^2_{\text{m}} = \frac{\chi^2_{\text{on,m}}}{N_{\text{on,m}}} + \frac{\chi^2_{\text{1xMM,m}}}{N_{\text{1xMM,m}}} + \frac{\chi^2_{\text{2xMM,m}}}{N_{\text{2xMM,m}}}, \quad \text{m} = \log_{10} k_{\text{clv}}, \log_{10} K_{\text{A}}. \tag{S11}$$

with $N_{\text{on,m}} = 1$, $N_{\text{1xMM,m}} = 20$ and $N_{\text{2xMM,m}} = 190$.

Using a sequence-independent model, we aggregate all data points sharing the same mismatch configuration in one weighted average value. If we measure value $\text{m}_{\text{i}}$, and have a measurement error of $\delta \text{m}_i$, we use the weighted average

$$\hat{\text{m}}_{[\text{mm-pattern}]} = \sum_{i \in \binom{\text{sequences with}}{\text{mm-pattern}}} w_i \text{m}_i \ , \quad w_i = \frac{1/\delta \text{m}_i^2}{\sum_{j \in \binom{\text{sequences with}}{\text{mm-pattern}}} 1/\delta \text{m}_j^2} \tag{S12}$$

to compare to our model prediction. The sums in **Equation** S12 run over all off-target sequences in the library that have the same mismatch pattern. This particular weighted average is chosen as it minimizes the square deviation if we have complete freedom in choosing the average for each position. Though our model is much more constrained then this, we use this weighing as a good fit to the weighted averaged data then has the potential to represents a good fit to the raw data. The error of the mean can be calculated as

$$\delta\hat{\mathrm{m}}^2_{[\text{mm-pattern}]} = \sum_{i\in\binom{\text{sequences with}}{\text{mm-pattern}}} w_i^2 \delta \mathrm{m}_i^2. \tag{S13}$$

Using the weighted averaged data points, the cost function is built up from

$$\chi^2_{\text{on,m}} = \frac{(\hat{\mathrm{m}}^{\text{model}}_{\text{on}} - \hat{\mathrm{m}}^{\text{experiment}}_{\text{on}})^2}{\delta\hat{\mathrm{m}}^2_{\text{on}}}, \quad \chi^2_{\text{1xMM,m}} = \sum_{n=1}^{20} \frac{(\hat{\mathrm{m}}^{\text{model}}_{[\text{n}]} - \hat{\mathrm{m}}^{\text{experiment}}_{[\text{n}]})^2}{\delta\hat{\mathrm{m}}^2_{[\text{n}]}}$$
$$\chi^2_{\text{2xMM,m}} = \sum_{n=1}^{19}\sum_{k=n+1}^{20} \frac{(\hat{\mathrm{m}}^{\text{model}}_{[\text{n,k}]} - \hat{\mathrm{m}}^{\text{experiment}}_{[\text{n,k}]})^2}{\delta\hat{\mathrm{m}}^2_{[\text{n,k}]}} \tag{S14}$$

The mismatch pattern $[n]$ represents a sequence with a single mismatch at position $n$, and $[n,k]$ represents a sequence with two mismatches, one at position $n$ and one at position $k$.

##### 3.2 Simulated-annealing optimization

The SA algorithm [3] is commonly used for high-dimensional optimization problems, such as the one presented here. The standard method is well established [3], and we here only give the special adjustments made to suit our problem. In every iteration we add numbers drawn from the uniform distribution on the interval $(-\delta, \delta)$ to each of the 41 model parameters representing free-energies and to the base 10 logarithm of those describing rates (see above). To initiate the fit, we fix $\delta$ (the step size) at 0.1 and adjust the effective temperature until an acceptance ratio between 40%-60%, based on the Metropolis condition, is reached. After this initial cycle, the effective temperatures follow an exponential cooling scheme with a 1% cooling rate ($T_{k+1} = T_k * 0.99^k$). At every temperature, we adjust the step size of the moves ($\delta$) until an acceptance ratio between 40-60% is reached, thereby enabling the parameters to change by smaller increments as the temperature decreases while avoiding accepting too many parameter configurations that worsen the $\chi^2$. This check is performed every 1000 steps. Once the check is passed, an additional 1000 steps are used to let the system 'equilibrate at temperature $T_k$' [3] before moving on the to next temperature ($k \to k+1$). As stop criteria we use

$$|\langle\chi^2\rangle_k - \langle\chi^2\rangle_{k+1}| \leq 10^{-5}\langle\chi^2\rangle_k \tag{S15}$$

In the above, $\langle\chi^2\rangle_k$ denotes the average of $\chi^2$ over the 1000 iterations at temperature $T_k$. To avoid premature stops, we further require the temperature to have dropped to one percent of its initial value.

To judge how reliably our fits find a minimum, we repeated our SA fit 55 times. Unfortunately, some fits clearly get stuck far from any global optimum, and in **Fig.** 2-3 we post-selected 30% of the best estimates as judged by our global cost function. These solutions differ on average by less than 2% in (log) association constants, and less than 9% in (log) cleavage rates. Though the discrepancy among some of the estimated parameters (**Fig.** 2) points to that we have not found the single global optimum, the fact that they all make essentially the same predictions points to that we have extracted what information we can from the data.

### 4 Removing undetermined quantities

**Fig.** 2 revealed that not all of our 44 parameters are tightly constrained by the given data. At the same time, some fit parameters are well determined, and we here coarse grain our model to remove the undetermined degrees of freedom.

#### 4.1 Local equilibration in metastable states

The on-target free-energy landscape shows a well-defined intermediate surrounded by two barriers of relatively high but poorly determined free energies. The rate of R-loop progression $k_\text{f}$ is also poorly determined. To understand why many fits still performed well, consider two facts: first, if a region is locally equilibrated, then the local distribution among states is independent of the forward rate $k_\text{f}$; second, if the time taken during the transition between locally equilibrated regions is negligible compared to the time spent in the locally equilibrated states before transitioning, then a change in $k_\text{f}$ can always be completely compensated for by a change in free energies in the barrier heights ($\Delta F$ ). To see the latter point, consider that the effective transition rate can be written as

$$k_\text{effective} \approx k_\text{f} e^{-\Delta F} \tag{S16}$$

To test the hypothesis that it is indeed a fundamental aspect of the process (local equilibration) that precludes us from finding a single optimum, we increased the on-target's free-energy values by a constant value of $\epsilon k_\text{B}T$, both for positions 1-8 and 13-8 (**Fig.** S2A). As a result, the two barriers surrounding the locally stable intermediate are increased from $\Delta F \to \Delta F + \epsilon$ (**Fig.** S2A). To compensate this barrier change we scale the R-loop progression rate as $k_\text{f} \to e^{\epsilon} \times k_\text{f}$ (**Fig.** S2B). **Fig.** S2C-D show that after rescaling the rate and barriers using $\epsilon = 2.5 k_B T$, the resulting prediction of NucleaSeq and CHAMP data does not change. We conclude that $k_\text{f}$ and $\Delta F$ are co-dependent due to local equilibration and rapid transitions between equilibrated states, which motivated us to construct the coarse-grained model as follows.

#### 4.2 The coarse-grained model

To construct the coarse-grained model put forth in **Fig.** 3, we start by recognizing a (locally) stable intermediate R-loop of length 11-12 nt in the on-target's free-energy landscape (**Fig.** 2A). Local equilibration means that the system equilibrates amongst the states 7-13 nt on a much shorter timescale than it can transition over the larger barriers in the free-energy landscape, from either the PAM bound open R-loop state (state 0) or the cleavage competent closed R-loop state (state 20).

The inverse average time to transition from the open to the intermediate state sets the effective transition rate $k_\text{OI}$. To calculate $k_\text{OI}$ we first determine the microscopic state with the minimum of free-energy between 7 and 13 denoted by location $n_\text{I}$. Next, we describe the evolution of the subsystem $\vec{P}_\text{OI}(t) = [P_0, P_1, P_2, ...., P_{n_\text{I}-1}]^T$ with the corresponding submatrix $K_{OI}$, that determines the evolution of our subsystem $\vec{P}_\text{OI}(t) = e^{-K_\text{OI} t}\vec{P}_\text{OI}(0)$ (**Equation** S4). As initial condition, all DNA are unbound.The probability distribution $\Psi_\text{OI}(\tau)$ of entry times into the intermediate state alone can be calculated as

$$\Psi_\text{OI}(\tau) = \frac{\partial P_{n_\text{I}(\tau)}}{\partial \tau} = -\sum_{n=0}^{n_\text{I}-1} \frac{\partial P_n(\tau)}{\partial \tau}, \tag{S17}$$

if we set $k_\text{b}(0) = 0$ and $k_\text{b}(n_\text{I}) = 0$. From here, we determine $k_\text{OI}$ as the (inverse) first moment of $\Psi_\text{OI}$,

$$\begin{aligned} k_\text{OI} &= \left(\int_0^\infty \tau \Psi_\text{OI}(\tau) \text{d}\tau\right)^{-1} = -\left(\sum_{n=0}^{n_\text{I}-1} \int_0^\infty \tau \frac{\partial P_n}{\partial \tau} \text{d}\tau\right)^{-1} \\ &= \left(\sum_{n=0}^{n_\text{I}-1} \int_0^\infty P_n(\tau) \text{d}\tau\right)^{-1} = \left(\sum_{n=0}^{n_\text{I}-1} \left(\int_0^\infty e^{-K_\text{OI}\tau} \text{d}\tau\right) P_n(0)\right)^{-1} \\ &= \left(\sum_{n=0}^{n_\text{I}-1} (K_\text{OI}^{-1} \vec{P}_\text{OI}(0))_n\right)^{-1}. \end{aligned} \tag{S18}$$

A similar process is used to calculate the rate of entering the closed state from the intermediate state, in which we construct the sub-system consisting of states $n_\text{I}$ through 19 and initiate the system at state

$n_{\rm I}$. The backward rates $k_{\rm IO}$ and $k_{\rm CI}$ can be determined in a similar manner, or directly by using the detailed balance condition (**Equation** S7), with coarse-grained energies

$$F_{\rm O} = F_{\rm PAM} = F_0, \quad F_{\rm C} = F_{20}, \quad F_{\rm I} = -\ln\left[\sum_{n=7}^{13} e^{-F_n}\right]. \tag{S19}$$

The last of these reflecting the free-energy of a system allowing to equilibrate between states 7 and 13. **Fig.** 3 shows the coarse-grained parameters, together with $k_{\rm on} = k_{\rm f}(-1)$, $k_{\rm off} = k_{\rm b}(0)$ and $k_{\rm clv} = k_{\rm f}(20)$ which keep their original interpretation.

#### Supplementary Figure Legends

**Supplementary Figure 1| Additional comparison to *in vitro* data (relates to Fig. 1).** **a**, Correlation plot of experimental cleavage rates for Cas9 (NucleaSeq) versus association constants (CHAMP). Both quantities are normalized and compared to their respective on-target value. **b-d**, Correlation plots of our model predictions (after fitting) versus (**b**) experimental cleavage rates from NucleaSeq, (**c**) association constants from CHAMP, and (**d**) binding rates from HiTS-FLIP. **e**, Association constants of CHAMP for library members with more than 2 mismatches. The library contained a series of mismatched off-targets, all containing different numbers of mismatches placed consecutively. Start- and end-points of these series of mismatches are indicated on vertical/horizontal axis. **f**, Validation of model; prediction of the association constants shown in **e**.

**Supplementary Figure 2| Co-dependency of microscopic model parameters (relates to Fig. 2).** **a**, Free-energy landscape obtained from our fits (blue) and a transformed landscape with the two barriers raised by 2.5 $k_{\mathrm{B}}T$ (pink). **b**, To compensate for the change in barrier height, we enhanced the rate of progressing the R-loop by a factor of $e^{2.5} \approx 12$ (**Supplementary Information**). **c**, Correlation of model predictions for association constants and **d**, cleavage rates for the original parameters vs adjusted parameters.

**Supplementary Figure 3| Coarse-grained model compared to complete model (relates to Fig. 3).** **a**, Model predictions for association constants from CHAMP using the complete 20-state kinetic model and the coarse-grained model described in **Fig. 3**. **b**, Same as **a**, for the cleavage rates from NucleaSeq.

**Supplementary Figure 4| additional precision-recall curves (relates to Fig. 5).** Precision-recall curves for our kinetic model (green), the CFD score (purple) and the uCRISPR score (orange) for the FANCF, VEGFA site 1, EMX1, HBB and RNF2 target sites. Curves are shown for all identified off-targets (union) and shared identified off-targets (intersection).

**Supplementary Figure 5| receiver-operator characteristic curves (relates to Fig. 5).** Receiver-operator characteristic curves for our kinetic model (green), the CFD score (purple) and the uCRISPR score (orange) for the FANCF, VEGFA site 1, EMX1, HBB and RNF2 target sites. Each curve displays the true positive rate (TPR) as a function of the false positive rate (FPR). Curves are shown for all identified off-targets (union) and shared identified off-targets ('intersection').

**Supplementary Figure 6 | F1 scores using the intersection of identified off-targets (relates to Fig 5).** F1-scores for our model (green), CFD prediction tool (purple) and uCRISPR (orange), for target sites EMX1, FANCF,HBB,RNF2 and VEGFA site 1. Off-targets shared across all experiments are included. For each condition, the maximum obtainable F1-score along the corresponding PR-curve is displayed (see **Supplementary Figure S4**).

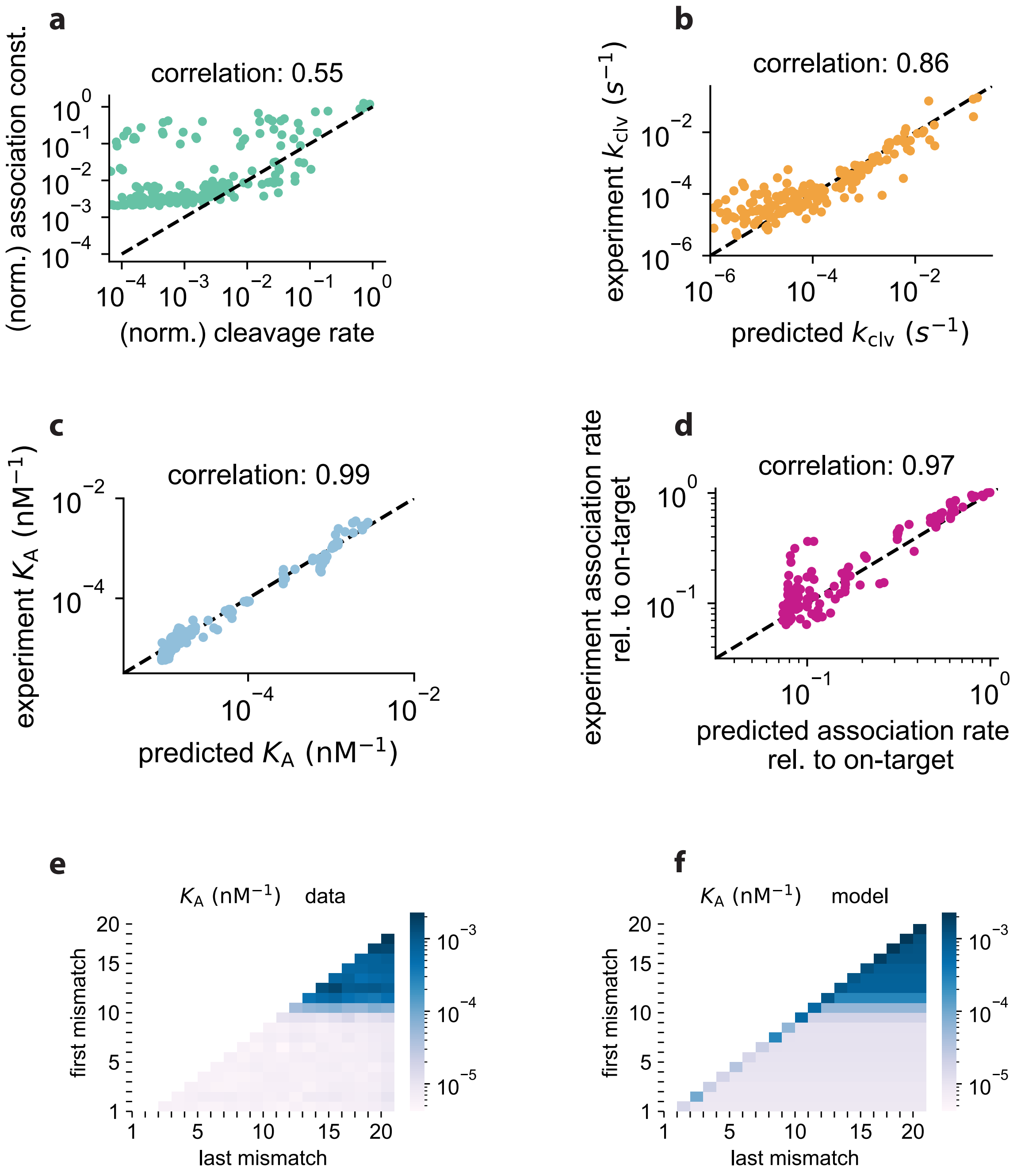


### Supplementary Figure 1

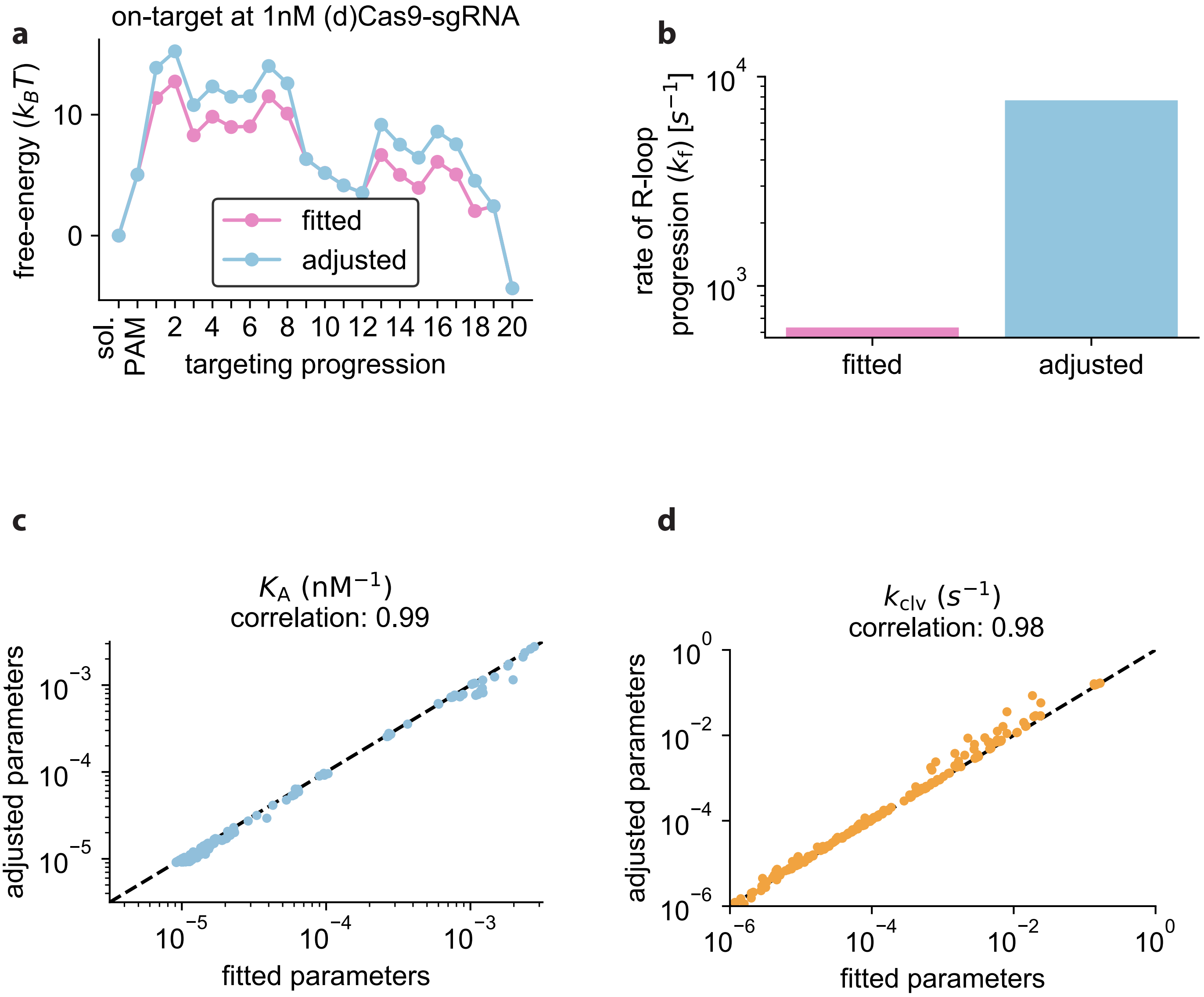


### Supplementary Figure 2

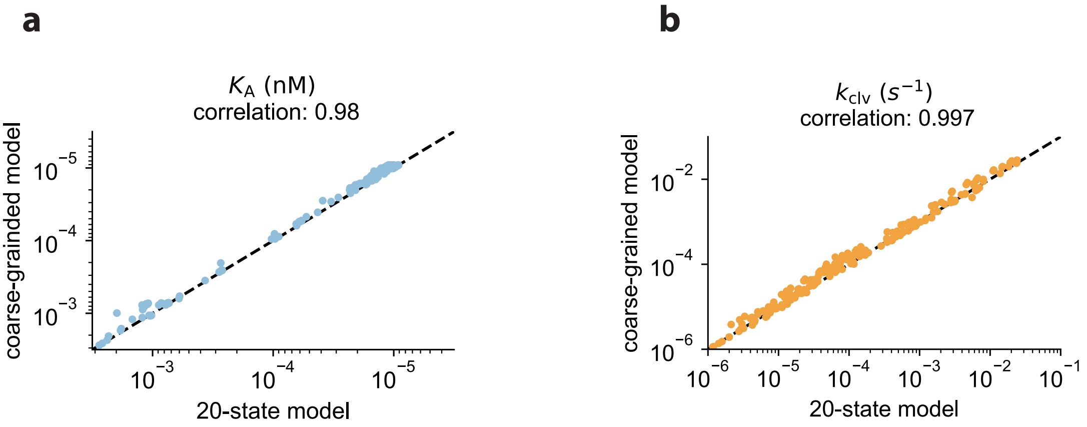


### Supplementary Figure 3

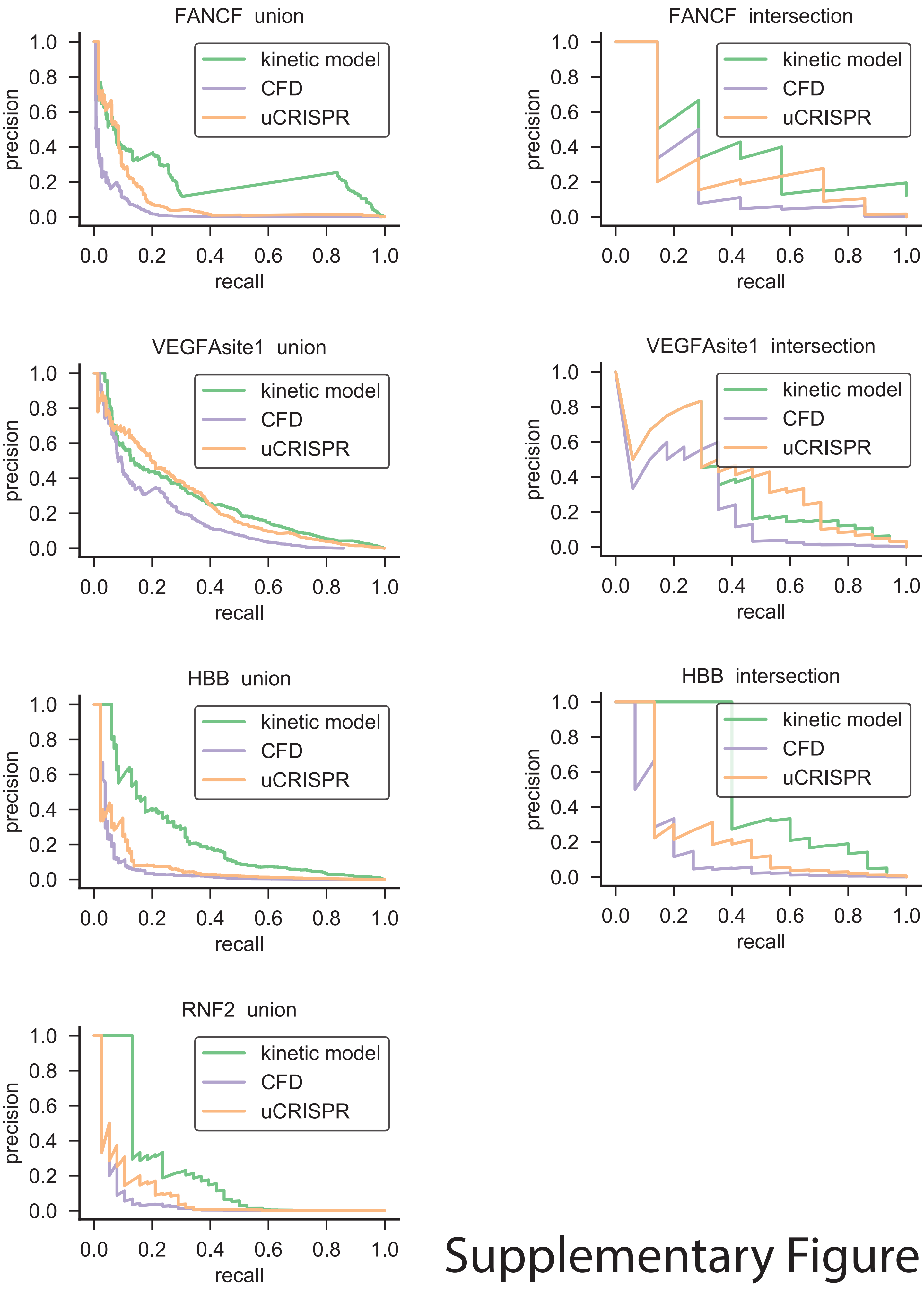


Supplementary Figure 4

### Supplementary Figure 5

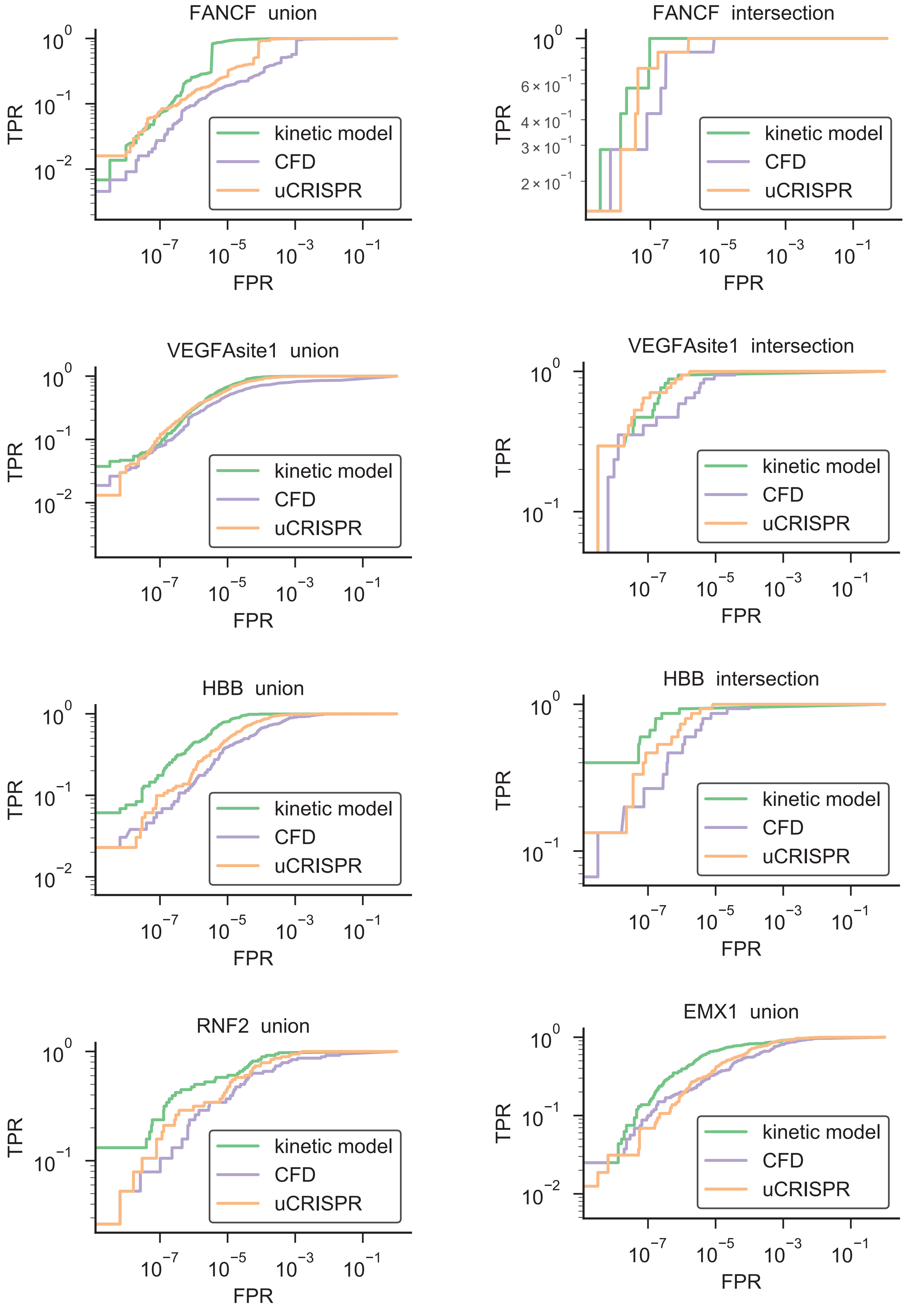

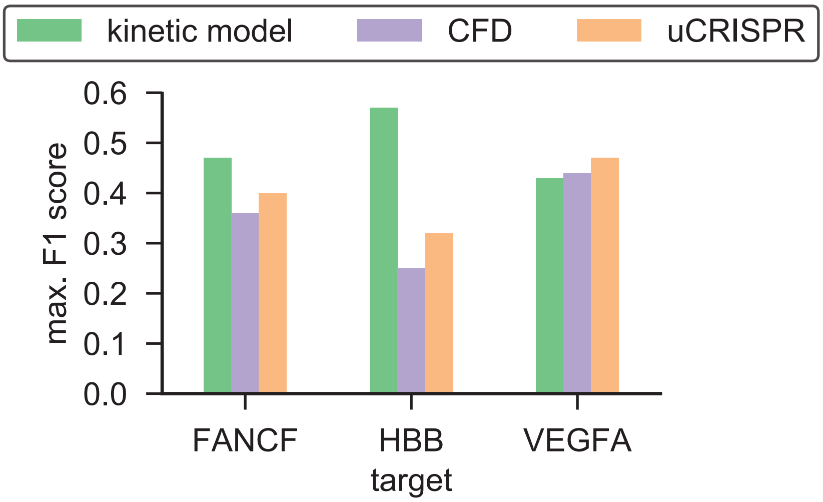


### Supplementary Figure 6
